# Information flows from hippocampus to auditory cortex during replay of verbal working memory items

**DOI:** 10.1101/2021.03.11.434989

**Authors:** Vasileios Dimakopoulos, Pierre Mégevand, Lennart Stieglitz, Lukas Imbach, Johannes Sarnthein

## Abstract

The maintenance of items in working memory relies on a widespread network of cortical areas and hippocampus where synchronization between electrophysiological recordings reflects functional coupling.

We investigated the direction of information flow between auditory cortex and hippocampus while participants heard and then mentally replayed strings of letters in working memory by activating their phonological loop. We recorded LFP from the hippocampus, reconstructed beamforming sources of scalp EEG, and - additionally in 3 participants – recorded from subdural cortical electrodes. When analyzing Granger causality, the information flow was from auditory cortex to hippocampus with a peak in the 4-8 Hz range while participants heard the letters. This flow was subsequently reversed during maintenance while participants maintained the letters in memory. The functional interaction between hippocampus and the cortex and the reversal of information flow provide a physiological basis for the encoding of memory items and their active replay during maintenance.

## 1 Introduction

Working memory (WM) describes our capacity to represent sensory input for prospective use [1, 2]. Maintaining content in WM requires communication within a widespread network of brain regions. The anatomical basis of WM was shown noninvasively with EEG / MEG [3-10] and invasively with intracranial local field potentials (LFP) [11-21] and single unit recordings [19, 21-24].

In cortical brain regions, WM maintenance correlates with sustained neuronal oscillations, most frequently reported in the theta-alpha range (4-12 Hz) [3-7, 9-20] or at even lower frequencies [25, 26]. Also in the hippocampus, WM maintenance was associated with sustained theta-alpha oscillations [15, 19]. As a hallmark for WM maintenance, persistent neuronal firing was reported during the absence of sensory input, indicating the involvement of the medial temporal lobe in WM [19, 22, 23].

At the network level, synchronized oscillations have been proposed as a mechanism for functional interactions between brain regions [27, 28]. It is thought that these oscillations show temporal coupling of the low-frequency phase for long-range communication between cortical areas [4, 6, 14, 17-19, 29]. This synchronization suggests an active maintenance process through reverberating signals between brain regions.

We here extend previous studies with the same task [3, 19] by recording from three participants with hippocampal LFP and cortical recordings (ECoG) from electrodes over primary auditory, parietal and occipital cortical areas. Given the low incidence of the epileptogenic zone in parietal cortex, parietal ECoG recordings are rare. In addition, we analyzed the directed functional coupling between hippocampal LFP and the beamforming sources of scalp EEG in all 15 participants. We found that the information flow was from auditory cortex to hippocampus during the encoding of WM items and the flow was from hippocampus to auditory cortex for the replay of the items during the maintenance period.

## 2 Results

### 2.1 Task and behavior

Fifteen participants (median age 29 y, range [18-56], 7 male, Table 1) performed a modified Sternberg WM task (71 sessions in total, 50 trials each). In the task, items were presented all at once rather than sequentially, thus separating the encoding period from the maintenance period. In each trial, the participant was instructed to memorize a set of 4, 6 or 8 letters presented for 2 s (encoding). The number of letters was thus specific for the memory workload. The participants read the letters themselves and heard them spoken at the same time. Since participants had difficulties reading 8 letters within the 2 s encoding period, also hearing the letters assured their good performance. After a delay (maintenance) period of 3 s, a probe letter prompted the participant to retrieve their memory (retrieval) and to indicate by button press (“IN” or “OUT”) whether or not the probe letter was a member of the letter set held in memory (**Fig. 1a**). During the maintenance period, participants rehearsed the verbal representation of the letter strings subvocally, i.e. mentally replayed the memory items. Participants had been instructed to employ this strategy and they confirmed after the sessions that they had indeed employed this strategy. This activation of the phonological loop [1] is a component of verbal WM as it serves to produce an appropriate behavioral response [2].

**Table 1:**
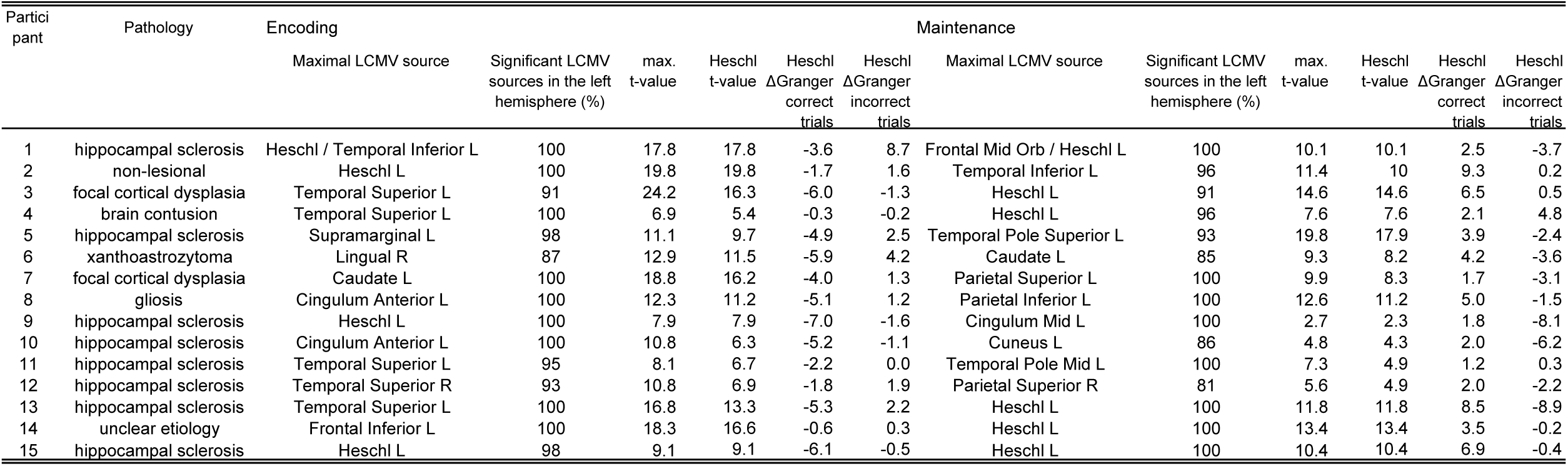
Participant characteristics and results of Granger causality analysis. For each participant we report the atlas parcels that contained EEG sources with the maximal t-value and the t-value of sources in auditory cortex (Heschl gyrus) during encoding and maintenance (non-parametric cluster-based permutation test p<0.05). In each participant, the vast majority of the significant LCMV sources was in the left hemisphere, both during encoding (≥ 87%) and during maintenance (≥ 81%) We also report the net information flow (ΔGranger) for correct and incorrect trials in the direction auditory cortex → hippocampus during encoding and in the direction hippocampus → auditory cortex during maintenance. **Abbreviations**: LCMV, Linearly constrained minimum variance; ΔGranger, difference of GC spectra.

**Figure 1.**
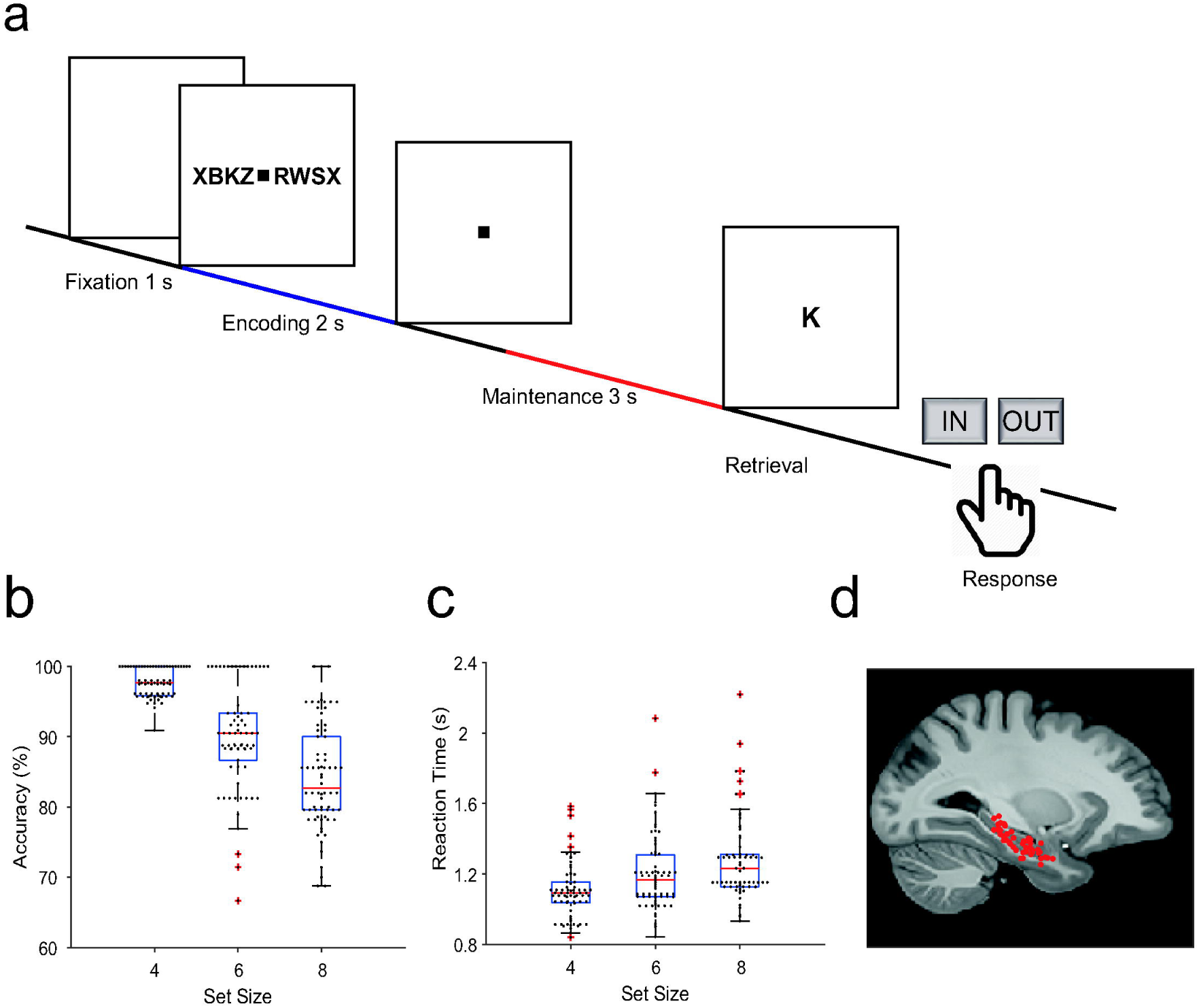
Task and recording sites. a) In the task, sets of consonants are presented and have to be memorized. The set size (4, 6 or 8 letters) determines WM workload. In each trial, presentation of a letter string (encoding period, 2 s) is followed by a delay (maintenance period, 3 s). After the delay, a probe letter is presented. Participants indicate whether the probe was in the letter string or not. b) Response accuracy decreases with set size (71 sessions). c) Reaction time increases with set size (53 ms/item). d) The tip locations of the hippocampal LFP electrodes for all participants (N = 15) are projected on the parasagittal plane x = 25.2 mm.

The mean correct response rate was 91% (both for IN and OUT trials). The rate of correct responses decreased with set size from a set size of 4 (97% correct responses) to set sizes of 6 (89%) and 8 (83%) (**Fig. 1 b**). Across the participants, the memory capacity averaged 6.1 (Cowan’s K, (correct IN rate + correct OUT rate - 1)*set size), which indicates that the participants were able to maintain at least 6 letters in memory. The mean response time (RT) for correct trials (3045 trials) was 1.1 ± 0.5 s and increased with workload from set size 4 (1.1 ± 0.5 s) to 6 (1.2 ± 0.5 s) and 8 (1.3 ± 0.6 s), 53 ms/item (**Fig. 1 c**). Correct IN/OUT decisions were made more rapidly than incorrect decisions (1.1 ± 0.5 versus 1.3 ± 0.6 seconds). These data show that the participants performed well in the task and that the difficulty of the trials increased with the number of letters in the set. In further analysis, we focused on correct trials with set size 6 and 8 letters to assure hippocampal activation and hippocampo-cortical interaction as shown earlier [19].

### 2.2 Power spectral density in cortical and hippocampal recordings

To investigate how cortical and hippocampal activity subserves WM processing, we analyzed the LFP recorded in the hippocampus (**Fig. 1 d**) together with ECoG from cortical strip electrodes (**Fig. 2 a, Fig. 3 a, f**). In the following, we present power spectral density (PSD) time-frequency maps from representative electrode contacts. In an occipital recording of Participant 1 (grid contact H3), strong gamma activity (> 40 Hz) in the relative power spectral density (PSD) occurred while the participant viewed the letters during encoding (increase >100 % with respect to fixation, **Fig. 2 b**). Similarly, encoding elicited gamma activity in a temporal recording over auditory cortex (increase >100%, grid contact C2, **Fig. 2 c**), similar as in [25]. Gamma increased significantly only in temporal and occipital-parietal contacts (permutation test with z-score > 1.96, **Fig. 2 a)**.

**Figure 2.**
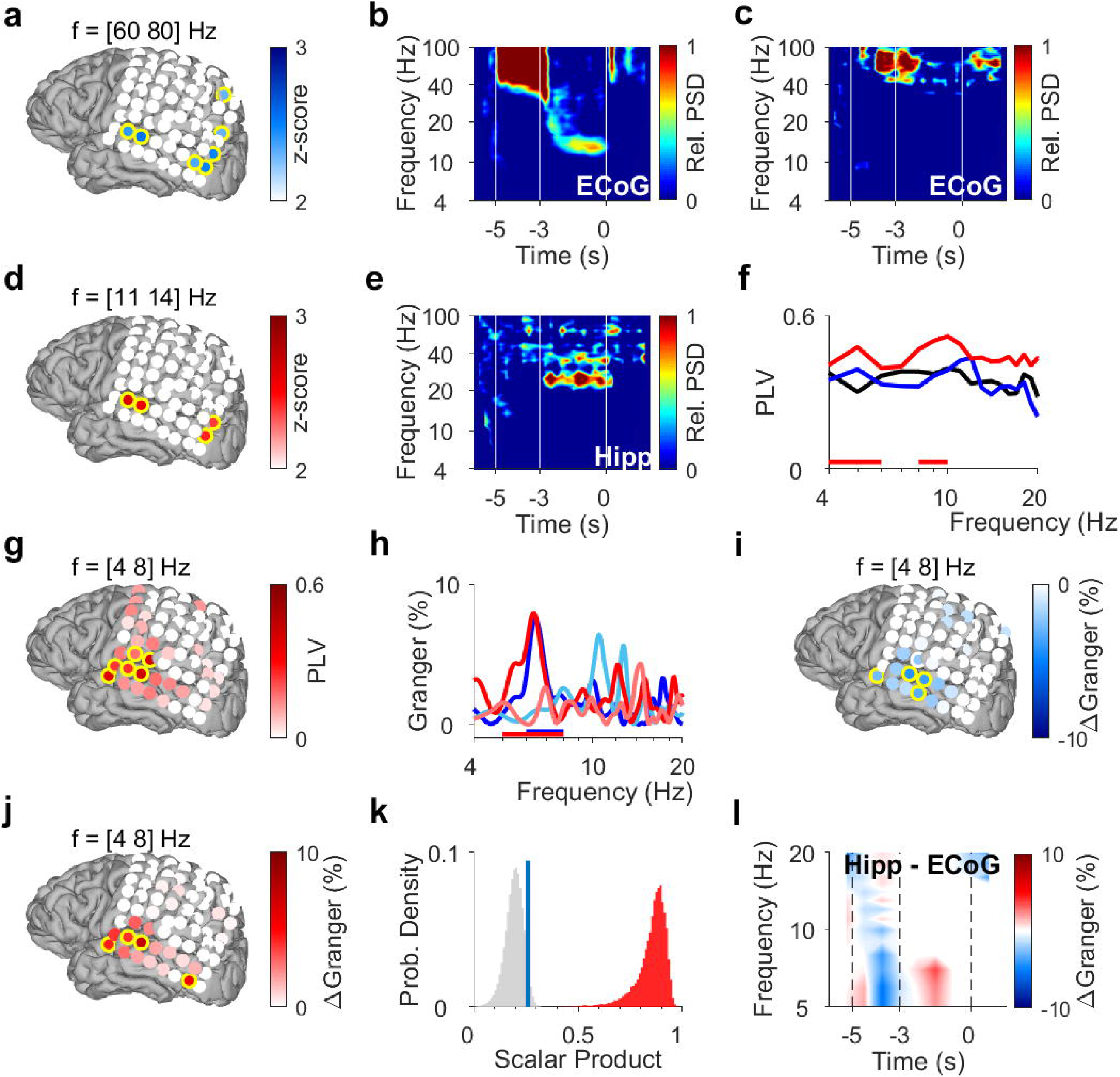
Encoding and replay of letters in Participant 1. a) Location of the ECoG contacts over temporal and parietal cortex for Participant 1. Relative gamma PSD ([60 80] Hz) during encoding ([-3.5 -3] s) is maximal for contacts over temporal and occipital-parietal cortex. b) The relative power spectral density (PSD) in the occipital contact (contact H3) over visual cortex shows gamma activity (>40 Hz) during encoding [-5 -3] s while the subject sees and hears the letters. Sustained low beta activity (11-14 Hz) appears towards the end of the maintenance period [-3 0] s. c) The relative power spectral density (PSD) in the temporal contact (contact C2) over auditory cortex shows gamma activity ([60 80] Hz) during the last second of encoding [-4 -3] s while the subject sees and hears the letters. d) Relative beta PSD ([11 14] Hz) during maintenance ([-2 0] s) is maximal for contacts over temporal and occipital cortex. e) Hippocampal PSD shows sustained beta activity towards the end of maintenance. f) Phase locking value (PLV) between hippocampus and auditory cortex (contact C3) during, fixation (black) encoding (blue) and maintenance (red). The PLV spectra show a broad frequency distribution. The PLV during maintenance is higher than during fixation. Red bars: frequency ranges of significant PLV difference (p<0.05, cluster-based nonparametric permutation test against a null distribution with scrambled trials during fixation and maintenance). g) Phase locking value (PLV) between hippocampus and cortex in theta (4-8 Hz) during maintenance ([-2 0] s) is highest to contacts over auditory cortex. h) Spectral Granger causality (GC). During encoding ([-5 -3] s), auditory cortex (contact C2) predicts hippocampus (6-8 Hz, dark blue curve exceeds light blue curve). During maintenance ([-2 0] s), hippocampus predicts auditory cortex (5-8 Hz, dark red curve exceeds light red curve). Bars: frequency range of significant ΔGranger (p<0.05), cluster-based nonparametric permutation test against a null distribution with scrambled trials during encoding (blue) and maintenance (red). i) Net information flow ΔGranger (4-8 Hz) during encoding ([-5 -3] s). ECoG over auditory cortex predicts hippocampal LFP. j) Net information flow ΔGranger (4-8 Hz) during maintenance ([-2 0] s). Hippocampus is maximal in predicting auditory cortex (contact C2 and surrounding contacts). k) Statistical significance of the spatial spread of contacts with high ΔGranger (4-8 Hz) during maintenance ([-2 0] s). We calculated the scalar product between two spread vectors. We then tested the statistical significance of the scalar product. The true distribution (red) is clearly distinct from the null distribution (gray, blue bar marks 95th percentile). l) The Granger time-frequency map illustrates the time-course of the spectra of panel h. During encoding, net information (ΔGranger) flows from auditory cortex to hippocampus (blue). During maintenance, the information flow is reversed from hippocampus to auditory cortex (red) indicating the replay of letters in memory. Grid contacts with significant increase are marked with a yellow rim (p<0.05, cluster-based nonparametric permutation test against a null distribution with scrambled trials). The time course in time-frequency maps is shown relative to the fixation period (b, c, e). To improve legibility, we present GC as Granger (%) = GC*100. Colors of Granger spectra indicate information flow: dark blue, cortex to hippocampus during encoding; light blue, hippocampus to cortex during encoding; dark red, hippocampus to cortex during maintenance; light red, cortex to hippocampus during maintenance. ΔGranger is the difference between spectra where ΔGranger < 0 denotes information flow cortex→hippocampus and ΔGranger > 0 denotes information flow hippocampus→cortex. Grid contacts are identified by column (anterior A to posterior H) and row (inferior 1 to superior 8).

**Figure 3.**
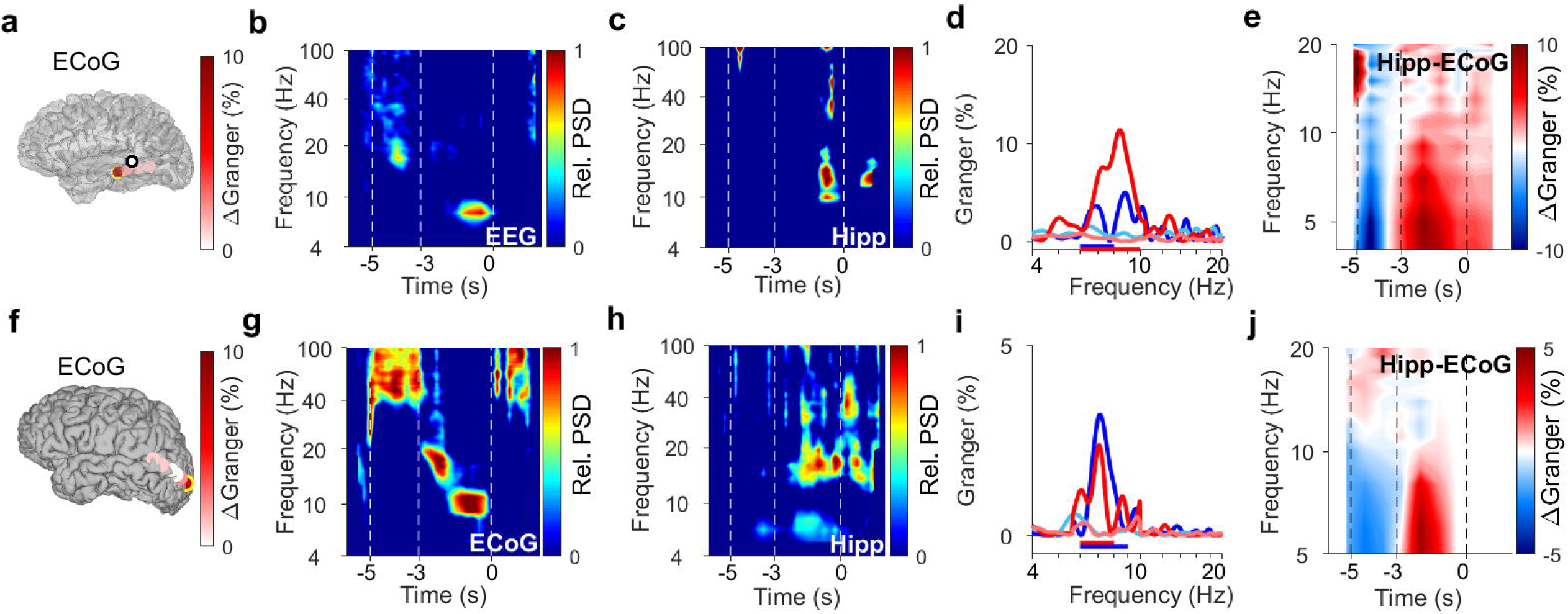
Encoding and replay of letters in two participants with ECoG. a) Location of the ECoG contacts in Participant 2. The most anterior strip contact records from auditory cortex. The black rimmed disk indicates the approximate location of scalp EEG electrode T5 over temporal cortex. Color bar: ΔGranger during maintenance (4-8 Hz). b) The relative power spectral density (PSD) in the temporal scalp EEG electrode (T5) shows beta activity (14-25 Hz) during encoding [-5 -3] s while the subject sees and hears the letters. Sustained theta activity (6-9 Hz) appears towards the end of the maintenance period [-3 0] s. c) Hippocampal PSD shows alpha-beta activity (9-18 Hz) towards the end of maintenance. d) Spectral Granger causality (GC). During encoding, the auditory cortex predicts hippocampus (6-8 Hz, dark blue curve exceeds light blue curve). During maintenance, hippocampal LFP predicts auditory cortex (6-10 Hz, dark red curve exceeds light red curve). e) The time-frequency map illustrates the time course of ΔGranger in Participant 2. f) Location of the ECoG contacts in Participant 3. The most posterior contact records from visual cortex (yellow rimmed disk). Color bar: ΔGranger during maintenance (4-8 Hz). g) The relative power spectral density (PSD) in the most posterior contact (yellow rimmed disk, panel f) shows gamma during encoding while the subject sees the letters. Sustained alpha activity (7-12 Hz) appears towards the end of the maintenance period. h) Hippocampal PSD shows sustained beta activity (13-21 Hz) towards the end of maintenance. i) Spectral Granger causality (GC). During encoding, the occipital ECoG predicts hippocampus (6-9 Hz, dark blue curve exceeds light blue curve). During maintenance, hippocampal LFP predicts ECoG (6-8 Hz, dark red curve exceeds light red curve). j) The time-frequency map illustrates the time course of ΔGranger in Participant 3. Grid contacts with significant increase are marked with a yellow rim (permutation test p<0.05). The time course in time-frequency maps is shown relative to the fixation period (b, c, g, h). To improve legibility, we present GC as Granger % = GC*100. Colors of Granger spectra indicate information flow: dark blue, cortex to hippocampus during encoding; light blue, hippocampus to cortex during encoding; dark red, hippocampus to cortex during maintenance; light red, cortex to hippocampus during maintenance. ΔGranger is the difference between spectra where ΔGranger < 0 denotes information flow cortex→hippocampus and ΔGranger > 0 denotes information flow hippocampus→cortex. Bars: frequency range of significant ΔGranger (p<0.05), cluster-based nonparametric permutation test against a null distribution with scrambled trials during encoding and maintenance, respectively.

After the letters disappeared from the screen, activity occurred in the low beta range (11-14 Hz, **Fig. 2 b**) towards the end of the maintenance period in temporal and occipital contacts (permutation test p < 0.05, **Fig. 2 d**). Similarly, the temporal scalp EEG of Participant 2 (black rimmed disk denotes electrode site T3 in **Fig 3 a**) showed activity during encoding and maintenance, albeit at lower frequencies (**Fig 3 b**); this pattern was found only in scalp EEG and not in ECoG, probably because the strip electrode was not located over auditory cortex. In Participant 3, a similar pattern occurred in the PSD of a temporo-parietal recording (most posterior strip electrode contact, **Fig 3 f**), where the appearance of the probe letter again prompted gamma activity (**Fig 3 g**). This site coincides with the generator of scalp EEG that was found in the parietal cortex for the same task [3]. The PSD thereby confirmed the findings of local synchronization of cortical activity during WM maintenance [3, 8, 9].

In the hippocampus of all three participants, we found elevated activity in the beta range (12-24 Hz) towards the end of the maintenance period (increase >100%, **Fig 2 e, Fig 3 c, h**), confirming the hippocampal contribution to processing of this task [19].

### 2.3 Functional coupling between hippocampus and cortex

To investigate the functional coupling between cortex and hippocampus, we first calculated the phase locking value (PLV). In Participant 1, we found high PLV over a broad frequency range in contacts over auditory cortex throughout the trial. Compared to encoding, maintenance showed enhanced PLV in the theta range between hippocampal LFP and cortical ECoG (PLV = 0.4 in contact C3, permutation test p<0.05, **Fig 2 f**). PLV in the [4-8] Hz theta range increased significantly with several contacts over auditory cortex (permutation test p<0.05, **Fig 2 g**). This speaks for a functional coupling between auditory cortex and hippocampus mediated by synchronized oscillations [26].

### 2.4 Directed functional coupling between hippocampus and ECoG

What was the directionality of the information flow during encoding and maintenance in a trial? We used spectral Granger causality (GC) as a measure of directed functional connectivity to determine the direction of the information flow between auditory cortex and hippocampus in Participant 1 during the trials. To improve legibility, we present GC as Granger (%) = GC*100. During encoding, the information flow was from auditory cortex to hippocampus with a maximum in the theta frequency range (dark blue curve in **Fig. 2 h**). The net information flow ΔGranger (GC hipp→cortex – GC cortex→hipp) during encoding was significant in the 6-8 Hz range (blue bar in **Fig. 2 h**, p<0.05 permutation test against a null distribution). During maintenance, the information flow in the theta frequency range was reversed (dark red curve), i.e. from hippocampus to auditory cortex (dark red curve in **Fig. 2 h**). The net information flow ΔGranger during maintenance was significant in the 5-8 Hz range (red bar in **Fig. 2 h**, p<0.05 permutation test against a null distribution). Concerning the spatial spread of the theta GC, the maximal net information flow ΔGranger (GC hipp→cortex – GC cortex→hipp) during encoding occurred from auditory cortex to hippocampus (p<0.05, permutation test, **Fig. 2 i**). During maintenance, the theta ΔGranger was significant from hippocampus to both auditory cortex and occipital cortex (permutation test p<0.05, **Fig. 2j**). Interestingly, in Participant 1, the distribution of high ΔGranger coincides with the distribution of high PLV: both show a spatial maximum to grid contacts over auditory cortex and both appear in the theta frequency range.

We next tested the statistical significance of the spatial spread of contacts with high ΔGranger (4-8 Hz) during maintenance ([-2 0] s). To provide a sound statistical basis, we tested the spatial distribution of GC on the grid contacts against a null distribution. The activation on grid contacts was reshaped into a grid vector. The spatial collinearity of two grid vectors was captured by their scalar product. We next performed 200 iterations of random trial permutations. For each iteration we selected two subsets of trials and we calculated the scalar product between the two vectors corresponding to these subsets. We then tested the statistical significance of the scalar product (**Fig. 2 k**). The true distribution (red) is clearly distinct from the null distribution (gray, blue bar marks the 95th percentile). The analogous procedure was performed for PSD (**Fig. 2 a, d**), PLV (**Fig. 2 g**) and GC during encoding (**Fig. 2 i**), which gave equally significant results in all cases.

As a further illustration of the ΔGranger time-course, the time-frequency plot (**Fig. 2 l**) shows the difference between GC spectra (GC hipp →cortex – GC cortex →hipp) at each time point, where blue indicates net flow from auditory cortex to hippocampus and red indicates net flow from hippocampus to auditory cortex.

Similarly in Participant 2, the time course of GC followed the same pattern between auditory cortex (anterior strip electrode contact in **Fig. 3 a**) and hippocampus (**Fig. 3 d,e**). Among the three participants that had both LFP and temporo-parietal ECoG recordings, Participant 3 had an electrode contact over visual cortex; the sensory localization was indexed by the strong gamma activity in the most posterior contact of the strip electrode (**Fig. 3 g**). The time-course of information flow between visual cortex and hippocampus (**Fig. 3 i,j**) followed the same pattern as described for the auditory cortex above. Thus, letters were encoded with information flow from sensory cortex to hippocampus; conversely, the information flow from hippocampus to sensory cortex indicated the replay of letters during maintenance.

### 2.5 Source reconstruction of the scalp EEG

We used beamforming [30] to reconstruct the EEG sources during encoding and maintenance for each of the 15 participants (Table 1). We tested whether the sources during fixation differed from sources during encoding and during maintenance (non-parametric cluster based permutation t-test [31, 32]). In each participant, the proportion of significant sources in the left hemisphere exceeded 80% of all significant sources. Across all participants, the spatial activity pattern during both encoding and maintenance showed the highest significance in frontal and temporal areas of the left hemisphere (**Fig S1**).

### 2.6 Directed functional coupling between hippocampus and averaged EEG sources

The synchronization between hippocampal LFP and EEG sources (N = 15 participants) confirmed the directed functional coupling found in the three participants with ECoG. We first calculated the GC between hippocampus and the EEG beamforming sources in the auditory cortex. We found that the mean GC spectra resembled the GC spectrum for ECoG in the theta frequency range ([4 8] Hz, **Fig. 4a**). During encoding, the net information flow was from auditory cortex to hippocampus (light blue curve – dark blue curve, blue bar, group cluster-based permutation test). During maintenance, the net information flow was reversed (dark red curve – light red curve, red bar, group cluster-based permutation test), i.e. from hippocampus to auditory cortex. Thus, both for ECoG and EEG sources, GC showed the same bidirectional effect in theta between auditory cortex and hippocampus.

**Figure 4.**
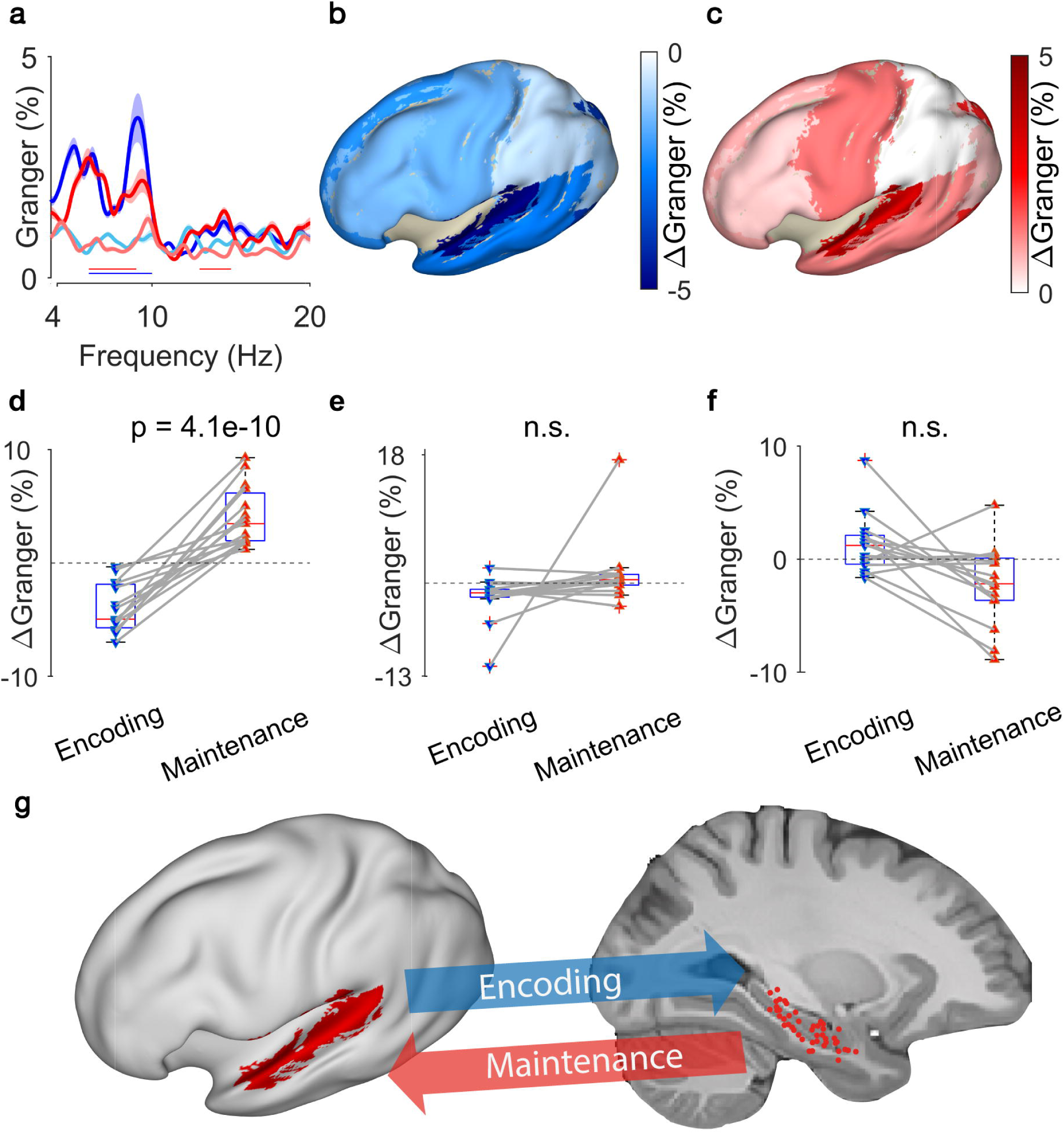
Granger causality between hippocampal LFP and EEG sources. a) Spectral Granger causality between hippocampal LFP and auditory EEG sources, averaged over all N=15 participants. The shaded area indicates the variability across the population. During encoding, the net Granger (ΔGranger) indicates information flow from auditory cortex to hippocampus ([6 10] Hz, blue bar). During maintenance, ΔGranger indicates information flow from hippocampal LFP to auditory cortex (red bars, [6 9] Hz, [13 15] Hz). Bars: frequency range of significant ΔGranger (p<0.05), group cluster-based nonparametric permutation t-test against a null distribution with scrambled trials during encoding and maintenance. Colors of Granger spectra indicate information flow: dark blue, cortex to hippocampus during encoding; light blue, hippocampus to cortex during encoding; dark red, hippocampus to cortex during maintenance; light red, cortex to hippocampus during maintenance. b) The median net information flow (ΔGranger) in the [4 8] Hz range during encoding is projected onto an inflated brain surface. The maximal ΔGranger appeared from temporal superior gyrus (median ΔGranger = -4.9%) indicating information flow from auditory cortex to hippocampus. Negative values of median ΔGranger appeared also in other areas, albeit less intense (ΔGranger > -3%). c) The median net information flow (ΔGranger) in the [4 8] Hz range during maintenance is projected onto an inflated brain surface. The maximal ΔGranger appeared from temporal superior gyrus (median ΔGranger = 3.4%) indicating an information flow from hippocampus to auditory cortex. Positive values of median ΔGranger appeared also in other areas, albeit less intense (ΔGranger < 2%). d) The maximal ΔGranger in the [4 8] Hz range was negative during encoding (blue, auditory cortex → hippocampus, median ΔGranger = -4.9%) and positive during maintenance (red, hippocampus → auditory cortex, median ΔGranger = 3.4%) for each participant (red and blue connected marker, paired permutation test, correct trials only). The mean values and statistical significance derive only from 10% of the correct trials in order to balance the number of incorrect trials. e) The net information flow between hippocampal LFP and lateral prefrontal cortex in the [4 8] Hz range has a lower median than to auditory cortex and higher variability (correct trials only, p = 0.16, paired permutation test, not significant). f) For incorrect trials, the maximal ΔGranger in the [4 8] Hz range is highly variable (p = 0.37, paired permutation test, not significant). g) Bidirectional information flow between posterior cortical sites and hippocampus in the working memory network. The Granger causality analysis suggest a surprisingly simple model of information flow during the task. During encoding, letter strings are verbalized as a melody; the incoming information flows from auditory cortex to hippocampus. During maintenance, participants actively recall and rehearse the melody in the phonological loop; Granger causality indicates an information flow from hippocampus to cortex as the physiological basis for the replay of the memory items.

To explore the spatial distribution, we computed GC also for other areas of cortex. We averaged the net information flow (ΔGranger) in the theta range across the participants and projected it onto the inflated brain surface (**Fig 4b, c**). During encoding, the mean information flow was strongest from auditory cortex to hippocampus (ΔGranger = -4.9 %, p = 0.0009, Kruskal-Wallis test, **Fig 4b**). For all other areas, the mean ΔGranger was also from cortex to hippocampus but the effect was weaker (mean ΔGranger = [-3 0]%, Dunn’s test, Bonferroni corrected). During maintenance (**Fig 4c**) the information flow was reversed. While all areas had information flow from hippocampus to cortex (ΔGranger = [0 2]%, Dunn’s test, Bonferroni corrected), the strongest flow appeared from hippocampus to auditory cortex (ΔGranger = 3.4%, p = 0.001, Kruskal-Wallis test).

### 2.7 Directed functional coupling and the participant’s performance

The reversal of ΔGranger appeared in all 15 participants individually (**Fig 4d**). We averaged ΔGranger for each participant in the [4-8] Hz theta frequency range. The ΔGranger between hippocampus and auditory cortex, was negative during encoding and was positive during maintenance in the (p = 4.1e-10, paired permutation test). The directionality and its reversal was missing for all other areas, e.g. lateral prefrontal cortex (p = 0.16, paired permutation test, **Fig 4e**). Of note, all analyses up to here were performed on correct trials only.

Finally, we established a link between the participants’ performance and ΔGranger. For incorrect trials, the net information flow ΔGranger from auditory cortex to hippocampus did not show the same directionality in all participants and did not reverse in direction (p = 0.37, paired permutation test, **Fig. 4f**). Since participants performed well (median performance 91%), we balanced the numbers of correct and incorrect trials. We calculated the GC in a subset of correct trials (median of 200 permutations of a number of correct trials that equals the mean percentage of incorrect trials = 10%); the effect was equally present for the subset of correct trials (p < 0.0005, **Fig 4d**). This suggests that timely information flow, as indexed by GC, is relevant for producing a correct response.

## 3 Discussion

Working memory (WM) describes our capacity to represent sensory input for prospective use. Our findings suggest that this cognitive function is subserved by bidirectional oscillatory interactions between the hippocampus and the auditory cortex as indicated by phase synchrony and Granger causality. In our verbal working memory task, the encoding of letter items is isolated from the maintenance period in which the active rehearsal of memory items is central to achieve correct performance. First, analysis of task-induced power showed sustained oscillatory activity in cortical and hippocampal sites during the maintenance period. Second, analysis of the inter-electrode phase synchrony and the directional information flow showed task-induced interactions in the theta band between cortical and hippocampal sites. Third, the directional information flow was from auditory cortex to hippocampus during encoding and, during maintenance, the reverse flow occurred from hippocampus to auditory cortex. This pattern was dominant on the left cortical hemisphere, as expected for a language related task. Fourth, the comparison between correct and incorrect trials suggests that the participants relied on timely information flow to produce a correct response. Our data suggests a surprisingly simple model of information flow within a network that involves sensory cortices and hippocampus (**Fig. 4 g**): During encoding, letter strings are verbalized as melody. The incoming information flows from sensory cortex to hippocampus (bottom-up). During maintenance, participants actively recall and rehearse the melody in their phonological loop [1, 2]. The Granger causality indicates the information flow from hippocampus to cortex (top-town) as the physiological basis for the replay of the memory items, which finally guides action.

The current study is embedded in previous studies using the same or similar tasks. Persistent firing of hippocampal neurons indicated hippocampal involvement in the maintenance of memory items [19, 22, 23]. An fMRI study reports salient activity in the auditory cortex during maintenance in an auditory working memory task [33], which indicates that sensory cortical areas are involved in the maintenance of WM items. During encoding, the activity of local assemblies was associated with gamma frequencies and local processing (**Fig. 2 a b c, Fig. 3 g**) while GC inter-areal interactions took place in theta frequencies, in line with previous reports [29, 34]. Parietal generators of theta-alpha EEG indicated involvement of parietal cortex in WM maintenance [3, 5, 7, 19, 35]. The hippocampo-cortical phase synchrony (PLV) was high during maintenance of the high workload trials [19]. Building on these previous studies, the current study focused on high workload trials and extended them by the analysis of directional information flow.

In the design of the task we aimed to separate in time the encoding of memory items from their maintenance. In the choice of the 2s duration for the encoding period were guided by the magic number 7±2, which may correspond to “how many items we can utter in 2 seconds” [1, 2]. The median Cowan’s k = 6.1 shows that high-workload trials were indeed demanding for the participants, where both encoding and maintenance may limit performance. We therefore presented the letters both as a visual and an auditory stimulus. Certainly, maintenance processes are likely to appear already during the encoding period as maintenance neurons ramp up their activity already during encoding [19]. Furthermore, encoding may extend past the visual stimulus (t = -3 s). We therefore focused our analysis on the last two seconds of maintenance [-2 0] s. With this task design, we found patterns of GC that were clearly distinct between encoding and maintenance.

The interaction between recordings from different brain regions has to be discussed with respect to volume conduction [36]. Of note, there was a strong frequency dependence of GC from hippocampus to ECoG **(Fig. 2 h, Fig. 3 d, i**). Likewise, GC to EEG sources showed a strong frequency dependence (**Fig. 4 a**). This speaks against volume conduction because the transfer of signal through tissue by volume conduction is independent of frequency in the range of interest here [37, 38]. Furthermore, there was a strong task dependence of GC (**Fig. 2 h, Fig. 3 d, i, Fig. 4 a**), again speaking against a strong contribution of volume conduction. Finally, the choice of two separate references for LFP and ECoG has been shown to avoid spurious effects in GC [39].

In the literature, there are several studies investigating the WM network. However, only few report directional interactions. One of these [17], reports cross-spectral directionality between intracranial recordings in frontal cortex and the medial temporal lobe in theta frequencies. One study on episodic memory suggests directional information flow to and from hippocampus [40]. Within hippocampus, directional information flow from posterior to anterior hippocampus indicated successful WM maintenance [20]. The frequencies of GC found in the current study were in the [4-8 Hz] theta band, in line with scalp EEG findings during WM tasks [4, 6] and other tasks [29] that activate oscillations in long-range recurrent connections [27, 28].

In sum, these results corroborated earlier findings on the working memory network and extended them by providing a physiological mechanism for the active replay of memory items.

## 4 Materials and Methods

### 4.1 Task

We used a modified Sternberg task in which the encoding of memory items and their maintenance were temporally separated (**Fig. 1a**). Each trial started with a fixation period ([−6, −5] s), followed by the stimulus ([−5, −3] s). The stimulus consisted of a set of eight consonants at the center of the screen. The middle four, six, or eight letters were the memory items, which determined the set size for the trial (4, 6, or 8 respectively). The outer positions were filled with “X,” which was never a memory item. The participants read the letters and heard them spoken at the same time. After the stimulus, the letters disappeared from the screen, and the maintenance interval started ([−3, 0] s). Since the auditory encoding may have extended beyond the 2 s period, we restrict our analysis to the last 2 s of the maintenance period ([−2, 0] s). A fixation square was shown throughout fixation, encoding, and maintenance. After maintenance, a probe was presented. The participants responded with a button press to indicate whether the probe was part of the stimulus. The participants were instructed to respond as rapidly as possible without making errors. After the response, the probe was turned off, and the participants received acoustic feedback regarding whether the response was correct or incorrect. The participants performed sessions of 50 trials in total, which lasted approximately 10 min each. Trials with different set sizes were presented in a random order, with the single exception that a trial with an incorrect response was always followed by a trial with a set size of 4. The task can be downloaded at www.neurobs.com/ex_files/expt_view?id=266.

### 4.2 Participants

The participants in the study were patients with drug resistant focal epilepsy. To investigate a potential surgical treatment of epilepsy, the patients were implanted with intracranial electrodes. The participants provided written informed consent for the study, which was approved by the institutional ethics review board (PB 2016-02055). The participants were right-handed and had normal or corrected-to-normal vision. For nine participants (4 – 13), the PSD and PLV has been reported in an earlier study [19].

### 4.3 Electrodes for LFP, ECoG, and EEG

The depth electrodes (1.3 mm diameter, 8 contacts of 1.6 mm length, spacing between contact centers 5 mm, ADTech®, Racine, WI, www.adtechmedical.com) were stereotactically implanted into the hippocampus. Subdural grids and strips were placed directly on the cortex according to the findings of the non-invasive presurgical evaluations. Platinum electrodes with 4 mm^2^ contact surface and 1 cm inter-electrode distances were used (ADTech®). In addition, scalp EEG electrodes were placed at the sites of the 10-20 system with minor adaptations to avoid surgical scalp lesions.

### 4.4 Electrode localization

To localize the ECoG grids and strips, we used the participants’ postoperative MR, aligned to CT and produced a 3D reconstruction of the participants’ pial brain surface. Grid and strip electrode coordinates were projected on the pial surface as described in [41] (**Fig. 2a, Fig. 3a,f**).

The stereotactic depth electrodes were localized using post-implantation computed tomography (CT) and post-implantation structural T1-weighted MRI scans. The CT scan was registered to the post-implantation scan as implemented in FieldTrip [42]. A fused image of CT and MRI scans was produced and the electrode contacts were marked visually. The hippocampal contact positions were projected on a parasagittal plane of MRI **(Fig. 1b**).

Some of the electrodes contacts were found in tissue that was deemed to be epileptogenic and that was later resected. Still, neurons in this tissue have been found to participate in task performance in an earlier study [19].

### 4.5 Recording setup, re-referencing, and preprocessing

All recordings were performed with Neuralynx ATLAS, sampling rate 4000 Hz, 0.5-1000 Hz passband (Neuralynx, Bozeman MT, USA, www.neuralynx.com). ECoG and LFP were recorded against a common intracranial reference. Signals were analyzed in Matlab (Mathworks, Natick MA, USA). We re-referenced the hippocampal LFP against the signal of a depth electrode contact in white matter. We re-referenced the cortical ECoG against a different depth electrode contact. The choice of two separate references for LFP and ECoG has been shown to avoid spurious functional connectivity estimates [39]. The scalp EEG was recorded against an electrode near the vertex and was then re-referenced to the averaged mastoid channels. All signals were downsampled to 500 Hz. All recordings were done at least 6 h from a seizure. Trials with large unitary artefacts in the scalp EEG were rejected. We focused on trials with high workload (set sizes 6 and 8) for further analysis. We used the FieldTrip toolbox for data processing and analysis [30].

### 4.6 Power spectral density

We first calculated the relative power spectral density (PSD) in the time-frequency domain (**Fig. 2 b**). Time-frequency maps for all trials were averaged. We used 3 multitapers with a window width of 10 cycles per frequency point, smoothed with 0.2 × frequency. We computed power in the frequency range 4-100 Hz with a time resolution of 0.1 s. The PSD during fixation ([−6.0, −5.0] s) served as a baseline for the baseline correction (PSD(t) – PSD(fixation))/ PSD(fixation) for each time-frequency point.

### 4.7 Phase locking value

To evaluate the functional connectivity of hippocampus and cortex, we calculated the phase-locking value (PLV) between hippocampal LFP channels and ECoG grid (multitaper frequency transformation with 2 tapers based on Fourier transform, frequency range 4-100 Hz with frequency resolution of 1 Hz).

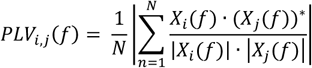

where PLVi,j is the PLV between channels i,j, N is the number of trials, X(f) is the Fourier transform of x(t), and (·)* represents the complex conjugate.

Using the spectra of the two-second epochs, phase differences were calculated for each electrode pair (i,j) to quantify the inter-electrode phase coupling. The phase difference between the two signals indexes the coherence between each electrode pair and is expressed as the PLV. The PLV ranges between 0 and 1, with values approaching 1 if the two signals show a constant phase relationship over all trials.

In our description of EEG frequency bands, we used theta (4-8 Hz), alpha (8-12 Hz), beta (12-24 Hz) and gamma (> 40 Hz), while the exact frequencies may differ in individual participants.

### 4.8 Source reconstruction of the EEG sources

We reconstructed the scalp EEG sources using linearly constrained minimum variance (LCMV) beamformers in the time domain. To solve the forward problem we used a precomputed head model template and aligned the EEG electrodes of each participant to the scalp compartment of the model. We then computed the source grid model and the leadfield matrix wherein we determined the grid locations according to the brain parcels of the automated anatomical atlas (AAL) [43]. We solved the inverse problem by scanning the grid locations using the LCMV filters separately for encoding and maintenance. The EEG sources were baselined with respect to the fixation period and presented as a percent of change from the pre-stimulus baseline. We defined cortical areas from multiple parcels since AAL is a parcellation based on sulci and gyri. We performed all the steps of the source reconstruction with FieldTrip [30] and projected the sources onto an inflated brain surface.

### 4.9 Spectral Granger causality

In order to evaluate the direction of information flow between the hippocampus and the cortex, we calculated spectral non-parametric Granger causality (GC) as a measure of directed functional connectivity analysis [30]. We evaluated the direction of information flow in the [4 20] Hz frequency range. To compute the GC we first downsampled the signals to the Nyquist frequency = 40 Hz. We then computed the GC between hippocampal contacts and ECoG grid contacts. We also computed GC between the same hippocampal contacts and EEG sources located over the regions of interest. GC examines if the activity on one channel can forecast activity in the target channel. In the spectral domain, GC measures the fraction of the total power that is contributed by the source to the target. We transformed signals to the frequency domain using the multitaper frequency transformation method (2 Hann tapers, frequency range 4 to 20 Hz with 20 seconds padding) to reduce spectral leakage and control the frequency smoothing.

We used a non-parametric spectral approach to measure the interaction in the channel pairs at a given interval time [44]. In this approach, the spectral transfer matrix is obtained from the Fourier transform of the data. We used the FieldTrip toolbox to factorize the transfer function H(f) and the noise covariance matrix Σ. The transfer function and the noise covariance matrix was then employed to calculate the total and the intrinsic power, S(f) = H(f)ΣH*(f), through which we calculated the Granger interaction in terms of power fractions contributed from the source to the target.

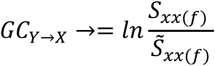

where S_(xx(f)) is the total power and 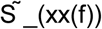 the instantaneous power. To improve legibility, we present GC as Granger % = GC*100. To average over the group of participants, we calculated the Granger spectra for the selected channel pairs and averaged these spectra over participants (**Fig 4a**).

To illustrate the time course of GC over time, we calculated time-frequency maps with the multitaper convolution method of Fieldtrip [30].

### 4.10 Statistics

To analyze statistical significance, we used cluster-based nonparametric permutation tests. To assess the significance of the difference of the Granger between different directions, we compared the difference of the true values to a null distribution of differences. We recomputed GC after switching directions randomly across trials, while keeping the trial numbers for both channels constant. Then we computed the difference of GC for the two conditions. We repeated this n = 200 times to create a null distribution of differences. The null distribution was exploited to calculate the percentile threshold p = 0.05. In this way, we compare the difference of the dark and light spectra against a null distribution of differences. We mark the frequency range of significant GC with a blue bar for encoding (dark blue spectrum exceeds light blue spectrum, information flow from cortex to hippocampus) and with a red bar for maintenance (dark red spectrum exceeds light red spectrum, information flow from hippocampus to cortex).

To test the statistical significance of the spatial spread of contacts with high PSD, PLV, or ΔGranger, we calculated the spatial collinearity on the grid contacts against a null distribution. First, we transform the activation on grid contacts into a grid vector. We then performed 200 iterations of random trial permutations. For each iteration we selected two subsets (50%) of trials and we calculated the scalar product between the vectors corresponding to the two subsets. The null distribution was created by randomly mixing trials from the two task periods fixation and encoding. We finally tested the statistical significance of the scalar product. The true distribution was established to be statistically distinct from the null distribution if it exceeded the 95th percentile of the null distribution.

We assess if the reconstructed EEG sources during encoding and maintenance are significantly different from the pre-stimulus baseline (fixation). We use the FieldTrip’s method ft_sourcestatistics [30] wherein we apply a non-parametric permutation approach to quantify the spatial activation pattern during the encoding of the memory items and their active replay.

Due to high average performance of the participants (91%) the number of correct and incorrect trials are imbalanced. To balance the number of correct trials with the number of incorrect trials we randomly selected 10% of the correct trials and recomputed the GC spectra and then the net information flow (ΔGranger). We repeated this n = 200 times and presented the mean ΔGranger for each participant.

For comparisons between two groups, we used the non-parametric paired cluster based permutation test. We created a null distribution by performing N = 200 random permutations.

To test the directionality of the information flow in the group of the participants, we used the group cluster based permutation t-test from the FieldTrip toolbox [30] with multiple comparison correction using the false discovery rate approach. Statistical significance was established at p < 0.05.

## Acknowledgments

We thank the physicians and the staff at Schweizerische Epilepsie-Klinik for their assistance and the patients for their participation. We acknowledge grants awarded by the Swiss National Science Foundation (SNSF 204651 to J. S.) and SNSF Ambizione fellowship (PZ00P3_167836 to P. M.) and a scholarship by Alexander S. Onassis Foundation (to V.D). The funders had no role in the design or analysis of the study.

## Ethical considerations

The participants provided written informed consent for the study, which was approved upfront by the institutional ethics review board (PB 2016-02055).

## Data and code availability

All data needed to evaluate the conclusions in the paper are present in the paper. All codes used to produce the results in the paper can be requested from the authors. The task can be downloaded at www.neurobs.com/ex_files/expt_view?id=266. Part of the data has been published earlier [35]. Additional data and code are indexed in www.hfozuri.ch.

## Author contributions

Conceptualization: JS; Methodology: VD, JS; Investigation: VD, JS, LS, LI; Visualization: VD, PM; Supervision: JS; Writing—original draft: JS, VD; Writing— review & editing: JS, VD

