## Supplementary Figure S1 for "Information flows from hippocampus to auditory cortex during replay of verbal working memory items"

**This PDF file includes:**

Fig. S1

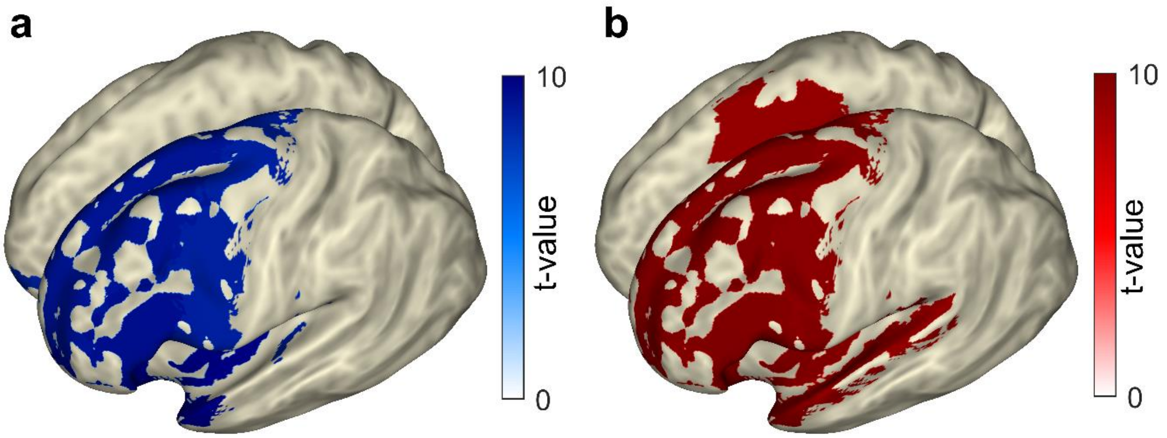


**Fig. S1. Spatial activation pattern of EEG beamforming sources**

a) The area of significant activation (t-value>8) during encoding compared to fixation is averaged for the group of participants and is projected onto an inflated brain surface. The most significant increase appears on sources over the left lateral prefrontal cortex. The spatial activation pattern at the cortical level spreads mostly over the left hemisphere (left frontal area, temporal pole, temporal superior gyrus and Heschl gyrus). On the right hemisphere, there is only a small orbitofrontal activation

b) The area of significant activation (t-value>8) during maintenance compared to fixation is projected onto an inflated brain surface. The most significant increase appears on sources over the left temporal superior gyrus (auditory cortex). The spatial activation pattern at the cortical level spreads mostly over the left hemisphere (left frontal area, temporal pole, temporal superior gyrus and Heschl gyrus). On the right hemisphere, an activation appears on premotor/motor cortex.

The spatial activation pattern derives from a non-parametric cluster based permutation t-test (N=1000 permutations, significance established at $t>1.96$ $p<0.05$). The activation map is thresholded at the 80% of the maximal t-value. Blue colorbar: encoding, red colorbar: maintenance.
